# Evidence for functional and non-functional classes of peptides translated from long non-coding RNAs

**DOI:** 10.1101/064915

**Authors:** Jorge Ruiz-Orera, Pol Verdaguer-Grau, José Luis Villanueva-Cañas, Xavier Messeguer, M Mar Albà

## Abstract

There is accumulating evidence that some genes have originated *de novo* from previously non-coding genomic sequences. However, the processes underlying *de novo* gene birth are still enigmatic. In particular, the appearance of a new functional protein seems highly improbable unless there is already a pool of neutrally evolving peptides that can at some point acquire new functions. Here we show for the first time that such peptides do not only exist but that they are prevalent among the translation products of mouse genes that lack homologues in rat and human. The data suggests that the translation of these peptides is due to the chance occurrence of open reading frames with a favorable codon composition. Our approach combines ribosome profiling experiments, proteomics data and non-synonymous and synonymous nucleotide polymorphism analysis. We propose that effectively neutral processes involving the expression of thousands of transcripts all the way down to proteins provide a basis for *de novo* gene evolution.

---

The mammalian genome is pervasively transcribed, this includes functional genes but also thousands of transcripts that are not conserved across species and which show weak or no signatures of natural selection^1–3^. Many of the latter transcripts are annotated as long non-coding RNAs (lncRNAs) because they lack conserved long open reading frames (ORFs). Recent studies based on the sequencing of ribosome-protected RNA fragments (ribosome profiling) have reported that a surprisingly large fraction of these transcripts is likely to translate small peptides^4–9^, although the significance of this finding has remained elusive.

Each ribosome profiling experiment generates millions of ribosome footprints that are subsequently mapped to the genome or the transcriptome to identify open reading frames (ORFs) that are being translated^10^. The codon-by-codon movement of the ribosome along the coding sequence results in a characteristic pattern of three nucleotide periodicity of the mapped reads, which makes ribosome profiling a very useful method to detect novel events of translation^4,11,12^. Given enough sequence coverage the technique can uncover low-abundant small peptides that would be otherwise difficult to detect by standard proteomics approaches^13,14^.

To assess the functional relevance of novel events of translation one can use the ratio between the number of non-synonymous and synonymous substitutions in the putative coding sequences^4,5^. However, this method requires an alignment of at least two homologous sequences. A more general approach that can be used in the absence of homology is the ratio between the number of non-synonymous and synonymous single nucleotide polymorphisms, compared to the one expected under neutrality. Under no selection, non-synonymous and synonymous polymorphisms accumulate at the same rate, whereas under purifying selection there is a deficit of non-synonymous polymorphisms because some amino acid changes disrupt the protein’s function^15^. Single nucleotide polymorphism analysis can be performed on a gene-by-gene basis or in pools of sequences that share certain features^2,16^.

We previously observed that, as a whole, putatively translated lncRNAs and young protein-coding genes share a number of similarities, such as small ORF size and weak selective constraints, compared with more widely conserved genes^8^. This pointed to a link between the translation of lncRNAs and the evolution of new proteins, but it did not solve the key question of whether translation of new ORFs could occur in the absence of selection at the protein level. This is a fundamental issue because for a new protein to acquire a function it first needs to be produced in the cell at significant amounts. Here by employing a combination of ribosome profiling data, sequence analysis and single nucleotide polymorphism information we obtain strong evidence that the majority of mouse proteins that are not conserved in rat or human selection evolve in a neutral manner. This study renders visible for the first time a layer of protein expression that is not dependent on selective processes, filling a gap in our understanding of the processes underlying *de novo* gene birth.

## Results

First we set to identify translated open reading frames (ORFs) in mouse protein-coding genes (codRNAs) and long non-coding RNAs (lncRNAs) using ribosome-profiling RNA-sequencing (Ribo-Seq) data from eight different tissues and cell lines (Supplementary Table 1 and references therein). In contrast to RNA sequencing (RNA-Seq) reads, which are expected to cover the complete transcript, Ribo-Seq reads correspond to regions bound by ribosomes. We mapped the RNA-Seq and Ribo-Seq reads to the mouse Ensembl gene annotations and, for the sake of completeness, also to a set of previously obtained novel mouse transcripts that did not correspond to annotated genes^3^.

We used the RibORF program^4^ to identify *bona fide* translated sequences among ORFs covered by at least 10 Ribo-Seq reads in transcripts expressed in one or more tissues (Fig. 1a and Supplementary Table 1). This program calculates a score for each ORF depending on the 3-nucleotide periodicity and uniformity of the mapped reads. Using a highly stringent RibORF score cut-off of 0.7^4^ we found that about 90% of the coding genes (15,020), and 20% of the annotated lncRNAs (539), were predicted to be translated in at least one sample. Additionally, we identified 286 genes that did not map to the gene annotations but contained translated ORFs (Fig. 1b). A widely used criterion to annotate a transcript as protein-coding is the presence of an ORF encoding a protein of at least 100 amino acids^17^. Not surprisingly, the vast majority of ORFs translated from annotated and novel lncRNAs encoded proteins smaller than 100 amino acids (smORFs).

**Figure 1.**
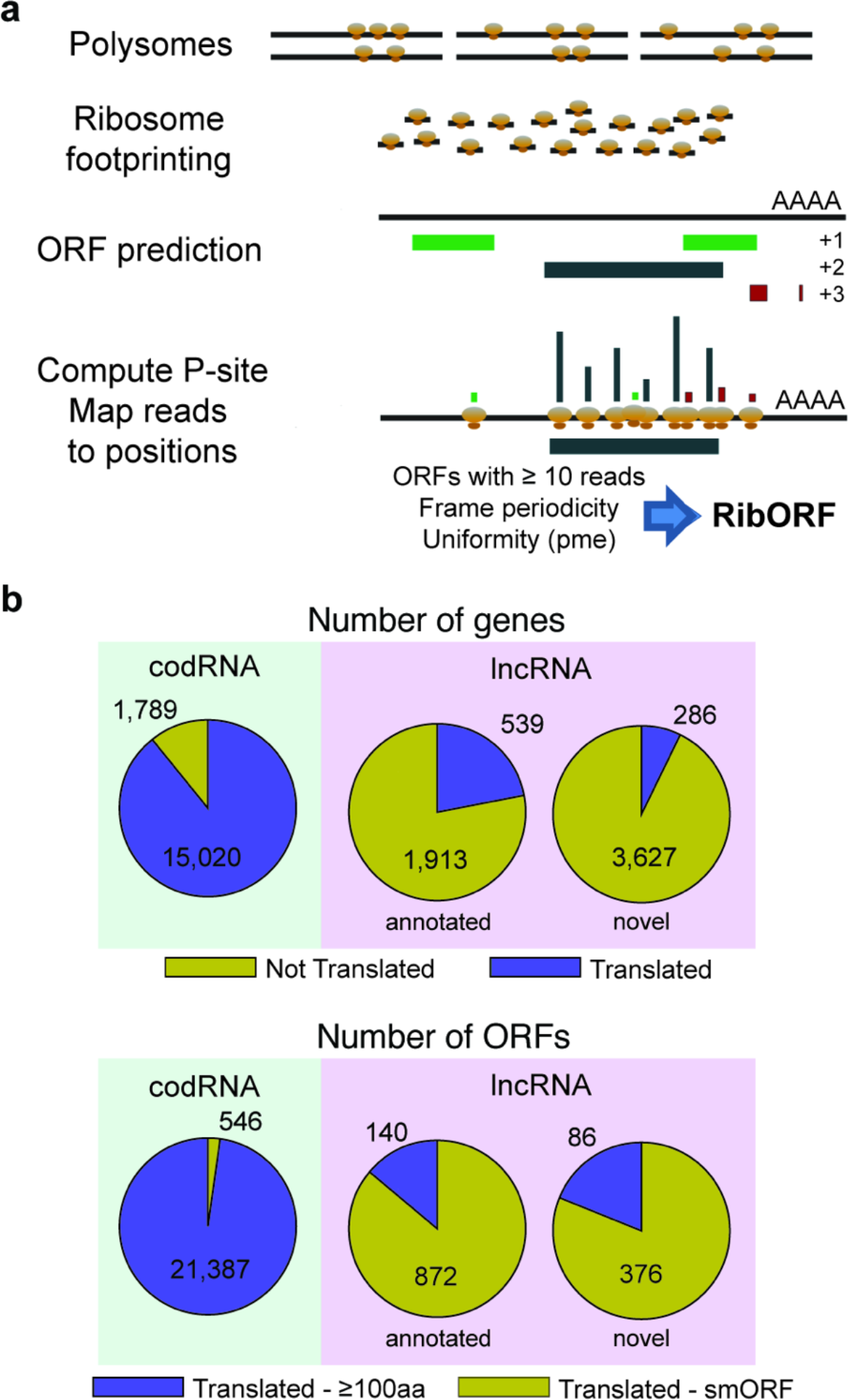
Detection of translated ORFs. **a.** Workflow to identify translated ORFs. Ribosome profiling (Ribo-Seq) reads, corresponding to ribosome-protected fragments, are mapped to all predicted canonical ORFs with length ≥ 30 nucleotides in transcripts. This is performed with single-nucleotide resolution after computing the read P-site per each read length. In each ORF, reads per frame and read uniformity are evaluated by RiboORF. **b.** Number of translated and non-translated expressed genes belonging to different classes after integrating data from eight different mouse tissues (Supplementary Table 1). **c.** Number of translated ORFs belonging to different classes. The translated ORFs have been divided into small ORFs (smORF, < 100 aa) and long ORFs (≥ 100 aa), depending on their length.

We hypothesized that some of the translated ORFs may evolve in a neutral manner and constitute a reservoir for the evolution of new protein-coding genes. To test this hypothesis, we first identified translated ORFs that were mouse-specific and then tested them for signatures of selection. We performed exhaustive sequence similarity searches of the ORFs against high coverage transcriptomes from human and rat as well as against the annotated proteomes of 101 different eukaryotic species (Fig. 2a, Supplementary Table 2 for a list of species, see Methods for more details). For these searches we discarded any proteins shorter than 24 amino acids, as the detection of homologues may be compromised in such cases due to lack of sufficient sequence information. We identified 1,980 different translated ORFs that showed no homology to expressed sequences in other species (class non-conserved or NC). In general, these ORFs had lower codon usage bias than conserved ORFs, as measured by a previously described hexamer-based coding score metric^8^ (Fig. 2b).

**Figure 2.**
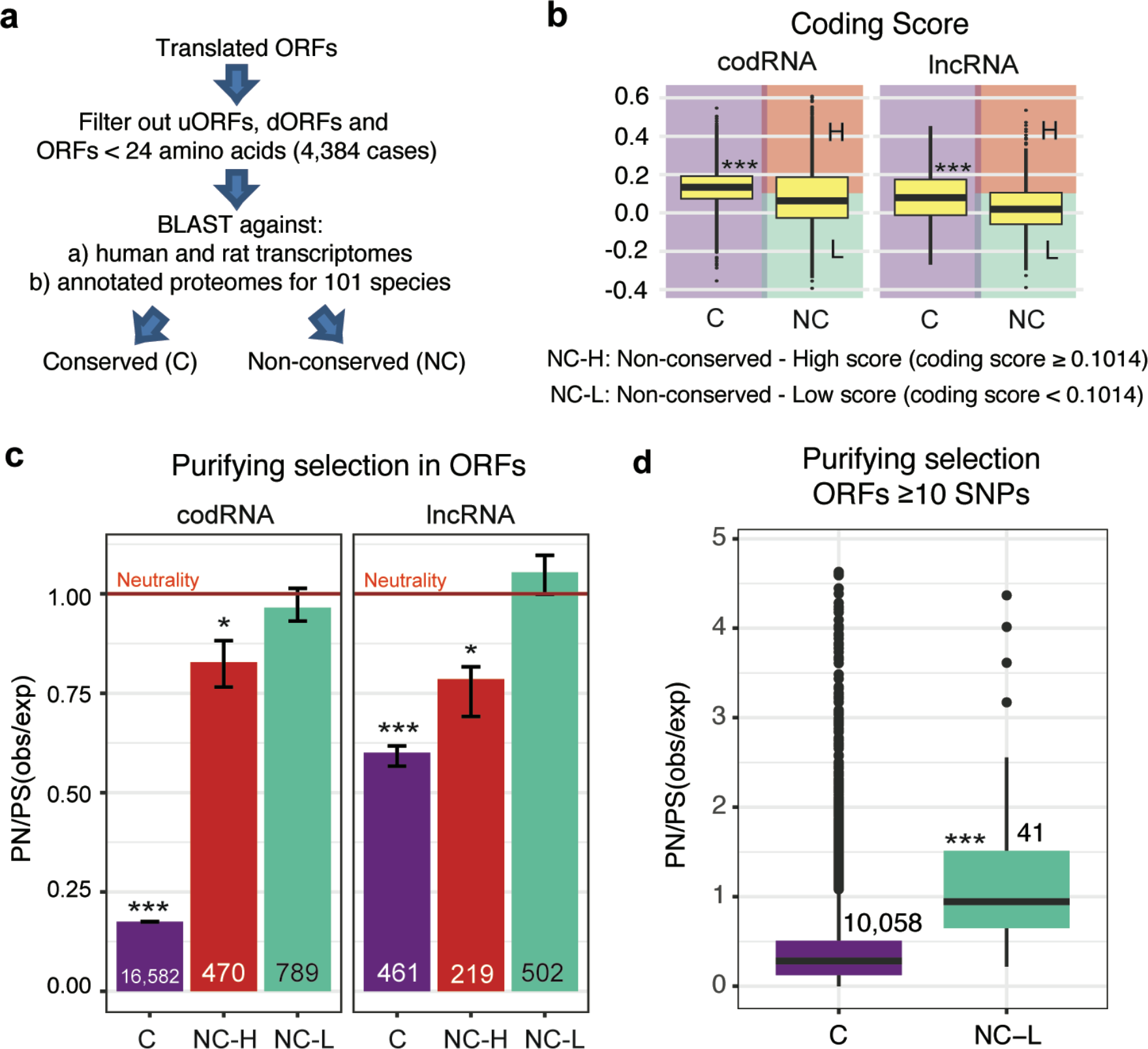
Identification of selection signatures. **a.** Workflow to identify conserved and non-conserved ORFs. Translated ORFs shorter than 24 amino acids, as well as non-conserved upstream and downstream ORF in conserved transcripts (uORFs and dORFs, see Methods), were filtered out. Any ORF with at least one BLAST match in another species was classified as conserved (C), otherwise it was classified as non-conserved (NC). **b.** Coding score in conserved (C) and non-conserved ORFs (NC). Conserved ORFs showed significantly higher coding score values than non-conserved ones; *** Wilcoxon test, p-value < 10^−5^. Non-conserved ORFs with a high coding score value (≥ 0.1014) were classified as NC-H, and the rest were classified as NC-L. **c.** Analysis of selective constraints in translated ORFs. PN/PS (obs/exp) refers to the normalized ratio between non-synonymous (PN) and synonymous (PS) single nucleotide polymorphisms; a value of 1 is expected in the absence of selection at the protein level. Conserved and NC-H ORFs showed significant purifying selection signatures. In contrast, NC-L ORFs did not show evidence of purifying selection at the protein level. Many conserved ORFs in lncRNAs are likely to encode functional micropeptides. Differences between observed and expected PN/PS were assessed with a chi-square test, * p-value < 0.05, *** p-value < 10^−5^. Error bars indicate the standard error of the sample proportion. Numbers of ORFs for the different categories are also displayed. **d.** Distribution of normalized PN/PS values for individual ORFs in different gene classes. Only ORFs with at least 10 SNPs were considered; the NC-H group contained too few cases to be analysed. The differences between C and NC-L are significant (Wilcoxon test, p-value <10^−5^).

To measure the strength of selection in conserved and non-conserved translated ORFs we employed a large collection of mouse single nucleotide polymorphisms (SNPs) for the house mouse subspecies *Mus musculus castaneus* ^18^. We could map a total of 324,729 SNPs to the set of translated ORFs. We grouped the ORFs into three different classes on the basis of conservation and coding score (Fig. 2b), and calculated the ratio between the number of observed non-synonymous and synonymous SNPs (PN/PS(obs)) in each class. We then normalized it by the same ratio expected under neutrality (PN/PS(exp)). The expected PN/PS was estimated using a table of nucleotide mutation frequencies in *Mus musculus castaneus* and the observed codon frequencies in each set of sequences of interest (Supplementary Tables 3 and 4). This allowed us not only to compare the strength of selection across different sets of sequences, as done in a previous study of ORFs translated from lncRNAs^8^, but also to discard selection if the normalized PN/PS was not significantly different from 1. Specifically, we used a chi-square test that compared the number of observed and expected non-synonymous and synonymous SNPs in each sequence set (Supplementary Table 5). As expected, the PN/PS of randomly selected ORFs from introns was approximately 1. Instead, the PN/PS of conserved ORFs was around 0.15 (Fig. 2c, chi-square test p-value < 10^−5^), consistent with protein functionality. One example in this group was Stannin^19,20^, a highly conserved peptide that regulates neuronal cell apoptosis (Fig. 3).

**Figure 3.**
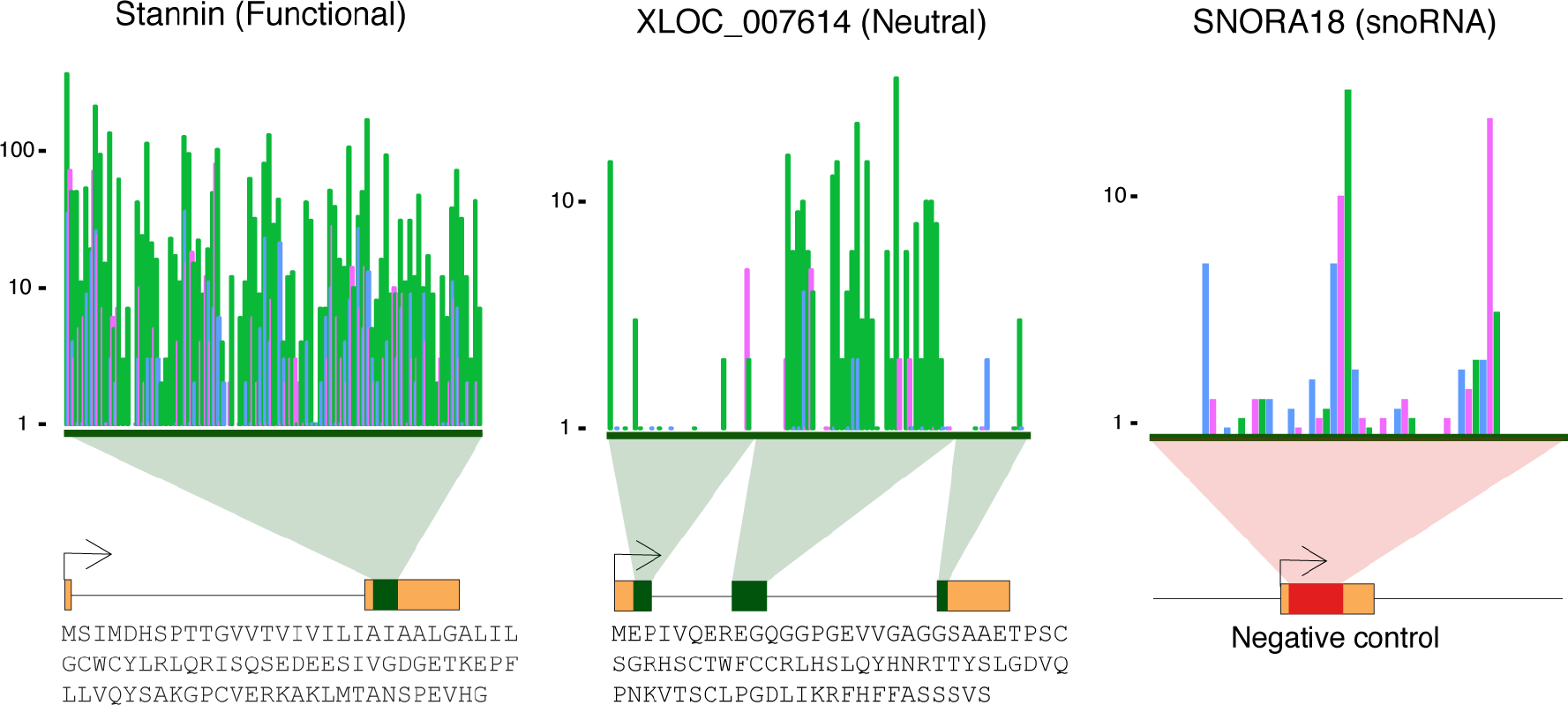
Three nucleotide periodicity of translated ORFs. The mapping of Ribo-Seq reads on different types of ORFs is shown. The Y axis represents the log-number of reads, the X axis the positions in the ORF. The reads show strong frame bias in the functional (conserved) and the neutral (NC-L) examples, with a preponderance of in-frame reads (green) versus off-frame reads (red and blue), while the frame bias is randomly distributed in the negative control (SNORA18). The exon/intron structure and the amino acid sequence for translated ORFs is also shown.

Non-conserved ORFs with high coding scores (NC-H coding score ≥ 0.1014, Fig. 2b and c, Supplementary Figure 1) had weak but significant signatures of selection (p-value < 0.05), possibly because of the existence of some functional mouse-specific genes. In contrast, the PN/PS ratio of the remaining non-conserved ORFs was not significantly different from 1, consistent with neutral evolution. Very similar results were obtained for non-conserved genes annotated as coding or lncRNA (Fig. 2c) and the two sets were merged into a single group of neutrally evolving ORFs (neutral ORFs). This set comprised about two thirds of the non-conserved ORFs (1,291 out of 1,980 ORFs analysed), and represented ~6.8% of the total number of mouse translated ORFs.

We used proteomics data from the PRIDE database^21^ to further validate the translation of this latter group of proteins. Despite their small size (median 44 amino acids), a limiting factor for their detection by standard proteomics-based tecniques^22^, we found proteomics evidence for 32 of the neutral ORFs (see Methods). This represents 2.5% of the proteins in this set (compared to less than 0.2% false positive rate, see Methods). This fraction is similar to the one obtained for conserved proteins subsampled to have a similar size distribution as the neutral ORFs (2.9%; in contrast, about 41% of all conserved ORFs have proteomics evidence). The test based on the PN/PS ratio confirmed that this subset of 32 ORFs did not deviate significantly from neutrality either (Supplementary Table 5).

The above analyses grouped the sequences into classes before computing the PN/PS ratio. In general, ORF-by-ORF analysis was not possible because the ORFs were small and contained too few SNPs. Nevertheless, 41 of the neutrally evolving ORFs contained 10 or more SNPs, and we decided to compute a normalized PN/PS ratio for these individual cases. The median PN/PS of these ORFs was around 1 and the distribution of PN/PS values was very different from that of conserved ORFs (Fig. 2d, Wilcoxon test, p-value < 10^−5^), consistent with the previous results. Finally, we quantified the number of ORFs that contained SNPs that generated premature stop codons, truncating more than half of the ORF, in the set of neutrally evolving ORFs and in the set of conserved ORFs. In the first case we found 72 out of 1,282 ORFs that contained this type of mutation (5.6%) and in the second case 296 out of 16,892 ORFs (1.75%). Considering that neutral ORFs are in general much shorter than conserved ORFs (median protein size 44 *versus* 412 amino acids), and thus less likely to accumulate ORF-truncating mutations by chance alone, the data clearly indicates a strong excess of ORF-truncating SNPs in neutral ORFs with respect to conserved ORFs. These analyses further support that the selective pressures acting on both kinds of ORFs are very different.

We next inspected in more detail the ribosome profiling patterns of neutral ORFs with respect to the rest of translated ORFs (hereafter called “functional”). Genes with a recent origin are usually ^23,24,3^ expressed at lower levels than older genes^23,24,3^ so it was not surprising to observe that neutrally evolving ORFs were associated with a lower number of Ribo-Seq reads per base than the rest of translated ORFs (median 0.193 versus 0.474, respectively, Supplementary figure 2). Consistent with translation, read periodicity in both neutral and functional ORFs was much higher than the random expectation of 0.33 (median values 0.70 and 0.80, respectively; see examples in Figure 3). Importantly, the results were highly reproducible across tissues (Figure 4a for hippocampus and embryonic stem cells; Supplementary Figure 3 hippocampus and brain), a result we would not expect in the case of spurious ribosome profiling signals. In general, the RibORF score of the translated ORFs was positively related to the number of mapped Ribo-Seq reads (Spearman correlation R=0.408), and to the size of the ORFs (Spearman correlation R=0.193). When we controlled for these two parameters, neutral and functional ORFs had equivalent distributions of RibORF score, periodicity and uniformity values (Figure 4b).

**Figure 4.**
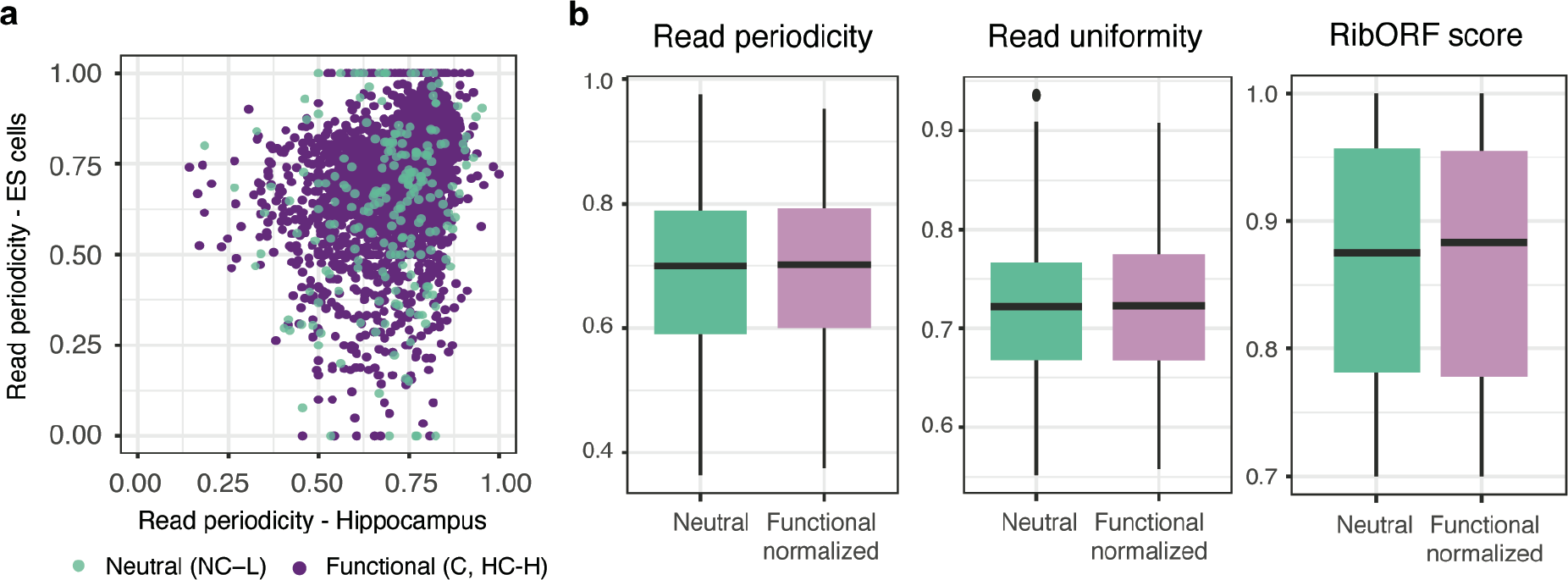
Properties of neutrally evolving ORFs. **a.** Relationship between the percentage of reads falling in the correct frame in neural embryonic stem cells cells and hippocampus samples, for neutral and functional ORFs having at least 10 reads in both samples and being translated in at least one sample. Spearman correlation coefficient is R=0.4224 for the neutrat set (p-value < 10^−5^) and R=0.4360 for the functional set (p-value < 10^−5^). **b.** Distribution of read periodicity, read uniformity and RibORF scores in neutral and functional translated ORFs after controlling for the number of Ribo-Seq reads and size of ORFs. The ‘functional normalized’ set is a randomly taken subset of the functional ORFs that has the same number of mapped Ribo-Seq reads and ORF size distribution as the set of neutrally evolving ORFs (n=900). Data is represented as box-plots for different number of read intervals; the box contains 50% of the data, horizontal line is the median value.

Subsequently, we compared our results to those obtained with two negative controls. The first control contained ORFs in alternative frames of annotated protein coding sequences with experimental protein evidence (“off-frame”). The second one contained randomly occurring ORFs in small nuclear and nucleolar annotated RNA sequences (“sRNA”). The latter RNAs are sometimes detected in ribosome profiling experiments due to the formation of ribonucleoprotein particles that protect the RNA from degradation^25^. As before, we only considered ORFs with at least 10 Ribo-Seq mapped reads. As expected, the vast majority of the ORFs in these controls did not display significant 3-nucleotide read periodicity (Supplementary figure 2, see a specific example in Figure 3). We found that only 234 out of 13,596 ORFs in “off-frame”, and 10 out of 304 ORFs in “sRNA”, had a RibORF score ≥0.7 (the threshold employed throughout our study). This corresponds to an overall false discovery rate (FDR) of 1.75%, much lower than the fraction of neutrally evolving proteins detected in our main analysis (6.8%).

Some transcripts contained relatively long ORFs but were not translated. One example of this sort was the previously described *de novo* non-coding gene *Poldi*^26^ that lacked any evidence of translation in the data we analysed. We next asked which factors may influence the translation of some neutrally evolving ORFs but not of others. First, we inspected the translation initiation sequence context but did not detect any significant differences between translated and non-translated ORFs (Supplementary Figure 4). We then hypothesized that the ORF coding score could affect the “translatability” of the transcript because codons that are abundant in coding sequences are expected to be more efficiently translated than other codons. Consistent with this hypothesis, we found that the translated neutrally evolving ORFs exhibited higher coding scores than non-translated ORFs with otherwise similar characteristics (Fig. 5a, Translated versus non-translated Wilcoxon test, p-value < 10^−5^). Importantly, we obtained a similar result after controlling for gene expression level (Fig. 5b, Wilcoxon test, p-value < 10^−5^). This is consistent with codon composition having an effect *per se* in ORF translation. When controlling by coding score, expression level, but not ORF length, had an effect on the translatability of the transcript (Fig. 5c).

**Figure 5.**
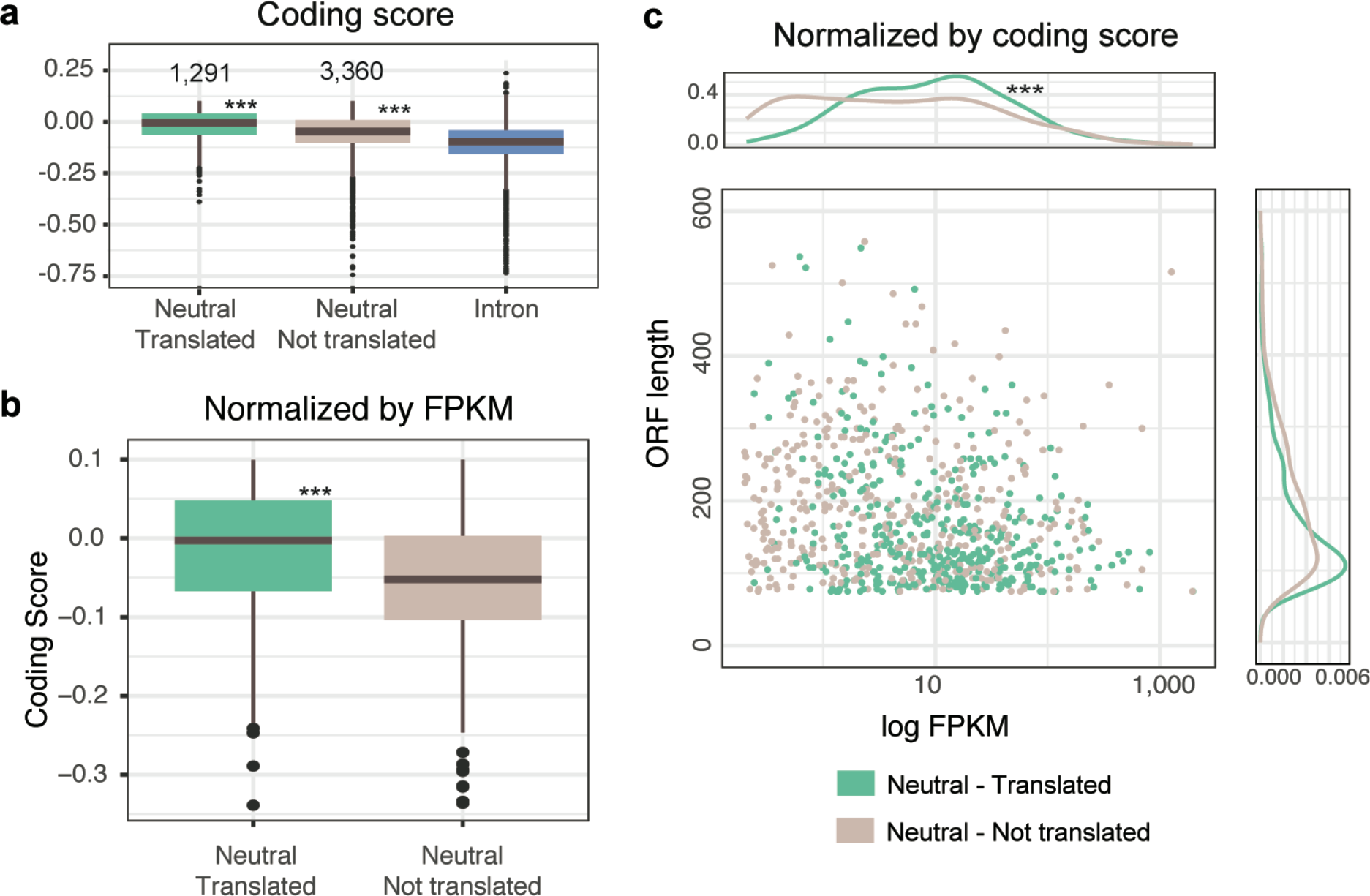
Factors influencing the translation of neutrally evolving ORFs. **a.** Influence of coding score in the translatability of neutrally evolving ORFs. Translated ORFs showed significantly higher coding score than non-translated ORFs, both sets had significantly higher coding scores than introns (Wilcoxon test p-value < 10^−5^, indicated by ***). **b.** Influence of coding score in the translatability of ORFs controlling for gene expression values, the two sets have comparable maximum FPKM gene expression (median FPKM value = 11.10). Translated ORFs showed significantly higher coding score values than non-translated ORFs; (Wilcoxon test p-value < 10^−5^). **c.** Influence of maximum FPKM gene expression and ORF length in the translatability of neutral ORFs normalized by coding score (median coding score value = −0.0052). Translated ORFs showed significantly higher FPKM values than non-translated ORFs (Wilcoxon test p-value < 10^−5^); differences in length were not significant.

The results suggest that the neutral ORFs that are translated are enriched in codons that are frequently found in functional protein coding sequences. This is consistent with the observation that abundant codons enhance translation elongation^27^, whereas rare codons might affect the stability of the mRNA^28^. It has been previously hypothesized that the distinction between translated and non-translated lncRNAs may be related to the relative amount of the lncRNA in the nucleus and the cytoplasm^4^. However, we found evidence that some lncRNAs with nuclear functions, such as *Malat1* and *Neat1,* were translated, suggesting that the cytosolic fraction of a transcript can be translated independently of its role or preferred location.

## DISCUSSION

The molecular mechanisms underlying *de novo* gene evolution are still poorly understood^29,30,31^. The sudden appearance of a new protein-coding gene from a genomic segment seems *a priori* highly improbable, but the process becomes much more likely if the genome is already being pervasively transcribed and translated outside functional protein-coding genes. An excess of transcription was already noted in the first large-scale cDNA sequencing efforts performed in human and mouse^32^, and more recent studies have found a high rate of transcriptional turnover when comparing closely related species^33^. Here we have shown that many of these transcripts are translated even if they only contain small ORFs, with the data currently available we have been able to identify 1,291 peptides in 1,132 genes that are likely to be of recent evolutionary origin and that show no signs of selection. This number is likely to be a gross underestimate because many transcripts are expressed at low levels, limiting their detection, and many cell types and tissues have not yet been sampled. According to recent estimates, the cost of transcription and translation in multicellular organisms is probably too small to overcome genetic drift^34^. Therefore, these activities may be effectively neutral. Our results indeed support that there is no barrier for the production of peptides that do not confer an immediate selective advantage.

The putative precursors of novel proteins identified here are of small size, which is consistent with observations for functional *de novo* genes identified in previous studies ^23,35–37^. We have also shown that random ORFs with a more favorable, coding-like, hexamer composition are more likely to be translated than other ORFs. Codon usage bias in functional sequences is related to the abundance of different tRNAs and correlates with expression level^38,39^. Thus, it seems logical that the translated neutral ORFs are biased towards those codons that are translated more efficiently.

The process of *de novo* gene origination involves the gain of a useful function by a previously non-functional sequence. The rate at which this happens remains to be determined but it has been observed that many random peptide sequences can function as secretion signals^40^, and selection for ATP-binding activity in a library of randomly generated 80 amino acid polypeptides successfully identified several candidates capable of binding to ATP^41^. Recent experiments performed in *E.coli* also suggest that random sequences can often affect cellular growth^42^. The pervasive translation of the transcriptome implies that *de novo* gene evolution has much more material at its disposal than previously thought.

## METHODS

### Transcript assembly

We used strand-specific polyA+ RNA sequencing data (RNA-Seq) data from different mouse and human tissues to assembly the species transcriptomes (Gene Expression Omnibus mouse GSE69241^3^, GSE43721^43^, and GSE43520^44^; human GSE69241^3^). The mouse RNA samples were extracted from strain Balb/C. RNA-Seq reads were filtered by length (> 25 nucleotides) and by quality using Condetri (v.2.2)^45^ with the following settings:-hq = 30 –lq = 10. We aligned the reads to the corresponding reference species genome with Tophat (v. 2.0.8, –N 3,-a 5 and –m 1)^46^. Multiple mapping to several locations in the genome was allowed unless otherwise stated. We assembled the transcriptome with Stringtie^47^, merging the reads from all the samples, with parameters-f 0.01, and-M 0.2. We used the species transcriptome as a guide (Ensembl v.75), including all annotated isoforms, but permitting the assembly of annotated and novel isoforms and genes (antisense, intergenic and intronic) as well. We excluded lncRNAs that overlapped annotated pseudogenes or that showed significant sequence similarity to known protein-coding sequences (BLASTP, e-value < 10^−4^). In the case of rat we employed a previously generated transcript assembly^48^.

### Ribosome profiling data

We used ribosome profiling data (Ribo-Seq) from 8 different mouse tissues or cell lines (see Supplementary Table 1), obtained from Gene Expression Omnibus under accession numbers GSE51424^49^, GSE50983^50^, GSE22001^51^, GSE62134^52^, GSE72064^53^, and GSE41246. Only datasets corresponding to non-pathogenic conditions were considered. The reads from the experimental replicates were merged before using RibORF to increase the resolution of the read periodicity, as done in the original RibORF paper^4^. For all analyses we considered only genes expressed at significant levels in at least one sample (RNA-Seq fragments per kilobase per Million mapped reads (FPKM) > 0.2). The expression of the genes detected in these samples is expected to be highly representative of the *Mus musculus* species as a whole. We mapped several brain RNA-Seq datasets from *Mus musculus castaneus*^33^ to the mouse assembled transcriptome using NextGenMap^54^. As expected, the vast majority of the genes expressed in brain samples from C57BL/6 mice^49^ also showed evidence of expression in *Mus musculus castaneus* brain RNA samples^33^(Supplementary Table 6).

We discarded anomalous reads (length < 26 or > 33 nt) and reads that mapped to annotated rRNAs and tRNAs in mouse from the Ribo-Seq sequencing datasets. Next, reads were mapped to the assembled mouse genome (mm10) with Bowtie (v. 0.12.7, parameters-k 1-m 20-n 1-best ‒‒strata). Considering that the ORFs had to be extensively covered by reads to be considered translated (high uniformity), we decided to include multiple mapped reads so as not to compromise the detection of paralogous proteins (Supplementary Fig. 7). We used the mapping of the Ribo-Seq reads to the complete set of annotated coding sequences in mouse to compute the position of the P-site (second binding site for tRNA in the ribosome) for reads of different size, as previously described^10,12^.

### Identification of translated ORFs

We predicted all translated ORFs (ATG to STOP) with a minimum length of 9 amino acids in the transcripts with RibORF (v.0.1)^4^. Only ORFs with a minimum of 10 mapped Ribo-Seq reads were considered. The RibORF classifier is based on a support vector machine algorithm, originally applied to human transcripts. The input parameters are the read periodicity and the read uniformity. The first one is the fraction of reads that correspond to the correct frame and the second one corresponds to the percentage of maximum entropy, a value of 1 indicates a completely even distribution of reads. For each ORF the program computes a score that depends on the values of these two parameters^4^. We used the same score cut-off as in the original paper (≥ 0.7), which had a reported false positive rate of 0.67% and false negative rate of 2.5%.

We eliminated any redundancy in the translated ORFs by taking the longest ORF when several overlapping translated ORFs were detected in the same gene. The identification of translated ORFs was done separately for the different tissues (Supplementary Table 1), and the data was subsequently integrated, taking the tissue with the highest RibORF score as representative. Differences in the number of translated ORFs in different tissues were related to the depth of sequencing and the number of reads that mapped to the top 5 most highly expressed proteins (Supplementary Fig. 6 and 7, respectively). For genes with no evidence of translation we selected the longest ORF across all transcripts for comparative purposes. Selecting the longest ORF was justified by the fact that, in translated ORFs, the ORF with the highest number of mapped Ribo-Seq reads was usually the longest ORF (75.7% for codRNAs and 84% for lncRNAs). We also generated a set of 4,013 randomly taken ORFs from introns, after discarding ORFs that showed significant sequence similarity to known proteins from the same species (BLASTP, e-value < 10^−4^).

We generated a negative control set by combining out-of-frame ORFs in mouse coding genes with experimental protein evidence according to Uniprot (“off-frame”) and randomly occurring ORFs in mouse small nuclear and nucleolar RNAs (“sRNAs”). These ORFs were required to have at least 10 Ribo-Seq mapped reads and were processed in the same manner as the main set of ORFs under study. The total number of sequences in the negative control was 13,900.

We also generated a positive control set composed of 2,163 randomly taken annotated mouse coding sequences with protein evidence in Uniprot. With these controls we estimated a false positive rate of 1.75% and a false negative rate of 2.54% for the above mentioned RibORF score cut-off.

### Sequence conservation

We searched for mouse translated ORF homologues in the human and rat transcriptomes using TBLASTN (limited to one strand, e-value < 10^−4^)^55^. We also performed sequence similarity searches against the annotated proteomes of 67 mammalian-species and 34 non-mammalian eukaryotes from a diverse range of groups compiled in a previous study^48^, using BLASTP (e-value < 10^−4^). For these searches we only considered query proteins of size 24 amino acids or longer, as shorter proteins may not contain sufficient information to perform homology searches. Mouse ORFs that did not have any homology hits in other species were classified as non-conserved, the rest as conserved. Translated non-conserved ORFs located upstream or downstream of another longer ORF in a conserved transcript (uORFs and dORFs) were excluded from this analysis.

We inspected the rat genomic syntenic regions of translated ORFs using LiftOver^56^. We classified the ORFs in two groups depending on whether the ORF was truncated in rat or not (the truncation had to affect more than half of the protein). For neutrally evolving ORFs the number of cases in which the ORF was truncated was similar to the number of cases in which it was not truncated, and in both cases the polymorphism patterns were consistent with neutrality (Supplementary Table 5). This indicated that, for this group, the presence of a similar ORF in rat does not imply functional conservation of the ORF. Therefore, we did not use information on rat genomic synteny to classify the genes as conserved/non-conserved.

### Single nucleotide polymorphism analysis

We obtained single nucleotide polymorphism (SNP) data from 20 individuals of the house mouse subspecies *Mus musculus castaneus^18^.* We classified SNPs in ORFs as non-synonymous (PN, amino acid altering) and synonymous (PS, not amino-acid altering). We calculated the PN/PS ratio in each ORF group by using the sum of PN and PS in all the sequences ((PN/PS)obs). We calculated the expected PN/PS under neutrality ((PN/PS)exp) using the mutation frequencies between pairs of nucleotides in *Mus musculus castaneus* and the codon composition of the different sequences or sets of sequences under study (Supplementary Tables 2 to 5). The observed transition to transversion ratio was 4.42, very similar to the 4.26 value obtained in early observations based on mouse-rat divergence data^57^. We tested for purifying selection by the number of observed and expected non-synonymous and synonymous SNPs using a chi-square test with one degree of freedom. Positively selected mutations are rapidly fixed in the population and their effect is expected to be negligible when using SNP data.

### Proteomics data

We used the proteomics database PRIDE^21^ to search for peptide matches in the proteins encoded by various gene sets. For a protein to have proteomics evidence, we required at least two distinct perfect matches of peptides that did not map to any other protein in the dataset, allowing for up to two mismatches. These are very stringent conditions with a false positive rate < 0.2%^48^.

### Coding score

We used a previously described metric based on hexamer frequencies to calculate the coding score of the sequences^8^. The method uses a table of pre-calculated hexamer scores that measure the relative frequency of each hexamer in coding versus non-coding sequences. These scores are then used to evaluate the coding propensity of a sequence based on its hexamer composition. The method has been implemented in a computational program called CIPHER that can be accessed online (http://evolutionarygenomics.upf.edu/cipher).

### Statistical tests and plots

The generation of plots and statistical tests was performed with the R package^58^.

### Data availability

Transcript assemblies from mouse, human and rat, as well as the mouse open reading frames (ORFs) predicted to be translated have been deposited at figshare (http://dx.doi.org/10.6084/m9.figshare.4702375). The code and executable file to calculate the coding score can be accessed at https://github.com/jorruior/CIPHER. The C program to calculated the PN/PS expected under neutrality is available at https://figshare.com/articles/computePNPS_c/5085706. Supplementary file 1 contains supplementary tables and figures.

## ACKNOWLEDGEMENTS

We are grateful for valuable discussions with many colleagues during this study. The work was funded by grants BFU2012-36820 and BFU2015-65235-P from Ministerio de Economía e Innovación (Spanish Government) and co-funded by FEDER (EC). We also received funding from Agència de Gestió d'Ajuts Universitaris i de Recerca Generalitat de Catalunya (AGAUR), grant number 2014SGR1121.

